# Gene fitness landscape of group A streptococcus during necrotizing myositis

**DOI:** 10.1101/432526

**Authors:** Luchang Zhu, Randall J. Olsen, Stephen B. Beres, Jesus M. Eraso, Matthew Ojeda Saavedra, Samantha L. Kubiak, Concepcion C. Cantu, Leslie Jenkins, Amelia R. L. Charbonneau, Andrew S. Waller, James M. Musser

## Abstract

Necrotizing fasciitis and myositis are devastating infections characterized by high mortality. Group A streptococcus (GAS) is a common cause of these infections, but the molecular pathogenesis is poorly understood. We report a genome-wide analysis using serotype M1 and M28 strains that identified novel GAS genes contributing to necrotizing myositis in nonhuman primates (NHP), a clinically relevant model. Using transposon directed insertion-site sequencing (TraDIS) we identified 126 and 116 GAS genes required for infection by serotype M1 and M28 organisms, respectively. For both M1 and M28 strains, more than 25% of the GAS genes required for necrotizing myositis encode known or putative transporters. Thirteen GAS transporters contributed to both M1 and M28 strain fitness in NHP myositis, including putative importers for amino acids, carbohydrates, and vitamins, and exporters for toxins, quorum sensing peptides, and uncharacterized molecules. Targeted deletion of genes encoding five transporters confirmed that each isogenic mutant strain was significantly impaired in causing necrotizing myositis in NHPs. qRT-PCR analysis showed that these five genes are expressed in infected NHP and human skeletal muscle. Certain substrate-binding lipoproteins of these transporters, such as Spy0271 and Spy1728, were previously documented to be surface-exposed, suggesting that our findings have translational research implications.

## Introduction

Necrotizing fasciitis, commonly known as the “flesh-eating disease”, is an invasive infection with very high rates of human morbidity and mortality (1, 2). In severe cases, contiguous muscle may be severely damaged, resulting in necrotizing myositis. Based on whether the cause is polymicrobial or monomicrobial, necrotizing fasciitis can be divided into type I (polymicrobial) and type II (monomicrobial) (2, 3). Group A streptococcus (GAS) is the primary cause of type II necrotizing fasciitis, and has an average case fatality rate of 29% (2, 4, 5). The molecular pathogenesis processes at work in GAS necrotizing fasciitis and myositis are poorly understood, a lack of knowledge that has impeded development of new effective diagnostics and therapeutics.

GAS is a human-specific pathogen causing more than 700 million infections annually worldwide (6). GAS infections range from relatively benign pharyngitis, skin, and soft-tissue infections, to life-threatening invasive diseases such as necrotizing fasciitis and necrotizing myositis (1, 7). No vaccine is currently available to prevent GAS infections. Decades of research have revealed some of the GAS molecules that contribute to the pathogenesis of necrotizing fasciitis and myositis, including M protein (8, 9), extracellular cysteine protease streptococcal pyrogenic exotoxin B (SpeB) (10–13), hyaluronic acid capsule (14), and cytotoxins NADase and streptolysin O (15–21). However, although the genome of GAS is relatively small (~1,800 genes) (22, 23), current understanding of the molecular pathogenesis of GAS necrotizing fasciitis and myositis is limited.

High-throughput genome-wide screens based on transposon mutagenesis strategies are very useful in providing new information about the genetic basis of bacterial virulence. Technologies such as signature-tagged mutagenesis (STM), transposon site hybridization (TraSH), and Tn-seq have been applied successfully to many bacterial pathogens to identify genes required for fitness under diverse *in vivo* and *ex vivo* conditions (24–30). In GAS, genome-wide transposon mutagenesis screens have been used to identify genes contributing to fitness during growth in human blood *ex vivo*, human saliva *ex vivo*, and mouse subcutaneous infections (24, 30–32). However, a genome-wide investigation of the GAS genes contributing to fitness in necrotizing myositis has not been undertaken.

Analysis of the molecular pathogenesis of GAS necrotizing myositis requires use of appropriate animal models. Toward this end, mouse and nonhuman primate (NHP) necrotizing myositis models have been developed that approximate this disease (10, 33, 34). Importantly, GAS is a human-specific pathogen. Some GAS virulence factors are specific for human and NHP target molecules (35–38) and have significantly decreased or no activity against the mouse homologs (38–40). Thus, NHP necrotizing myositis provides the most relevant experimental model possible.

Here we report the first use in NHPs of a relatively new genome-wide transposon mutagenesis technique termed transposon directed insertion-site sequencing (TraDIS) (41, 42). TraDIS was recently documented to provide much novel information about GAS genes contributing to fitness in human saliva (31). Using highly saturated transposon mutant libraries made in two genetically distinct GAS strains that are common causes of severe human infections, we identified novel genes required for bacterial fitness during necrotizing myositis. Our screen revealed the theme that GAS transporters play a pivotal role in this infection. We confirmed and extended the TraDIS screen data using isogenic mutant strains, *in vitro* growth phenotyping and qRT-PCR analysis of necrotic myositis tissues taken from infected NHPs and human patients.

## Results

### Construction of highly saturated transposon mutant libraries in genetically representative strains of serotype M1 and M28 of GAS

Transposon insertion mutant libraries were generated using serotype M1 strain MGAS2221 and serotype M28 strain MGAS27961 as the parental organisms. Strains of these two M protein serotypes were used because they are among the five more abundant M protein types causing invasive infections in many countries, and in some cases they are the dominant causes of infections. Thus, serotype M1 and M28 GAS are clinically highly relevant (43–46). These two strains were chosen for transposon mutagenesis because (i) strain MGAS2221 is genetically representative of a pandemic serotype M1 clone that arose in the 1980s, rapidly spread worldwide and currently is the most prevalent cause of severe infections globally (18, 20, 47), (ii) strain MGAS27961 is genetically representative of a virulent serotype M28 clone that is prevalent in the United States and elsewhere (48), (iii) both strains have wild-type alleles of all major transcriptional regulators that are known to affect GAS virulence, such as *covR* and *covS*, *ropB*, *mga*, and *rocA*, and (iv) both strains have been used previously in animal infection studies (18, 20, 33). Using transposon plasmid pGh9:IS*S1* (49), we generated dense transposon mutant libraries in strains MGAS2221 and MGAS27961 containing 154,666 (an insertion every 12 bp on average) and 330,477 (an insertion every 5.5 bp on average) unique transposon insertions, respectively (Figure 1A). This means that on average, the serotype M1 and M28 libraries had 66 and 139 insertions per open reading frame. The insertion index (number of unique insertions/size of the gene) of each of the genes in the M1 and M28 genomes is illustrated in Figure 1B. Use of the MGAS2221 transposon mutant library to identify novel genes contributing to GAS fitness in human saliva *ex vivo* has been described recently (31).

**Figure 1.**
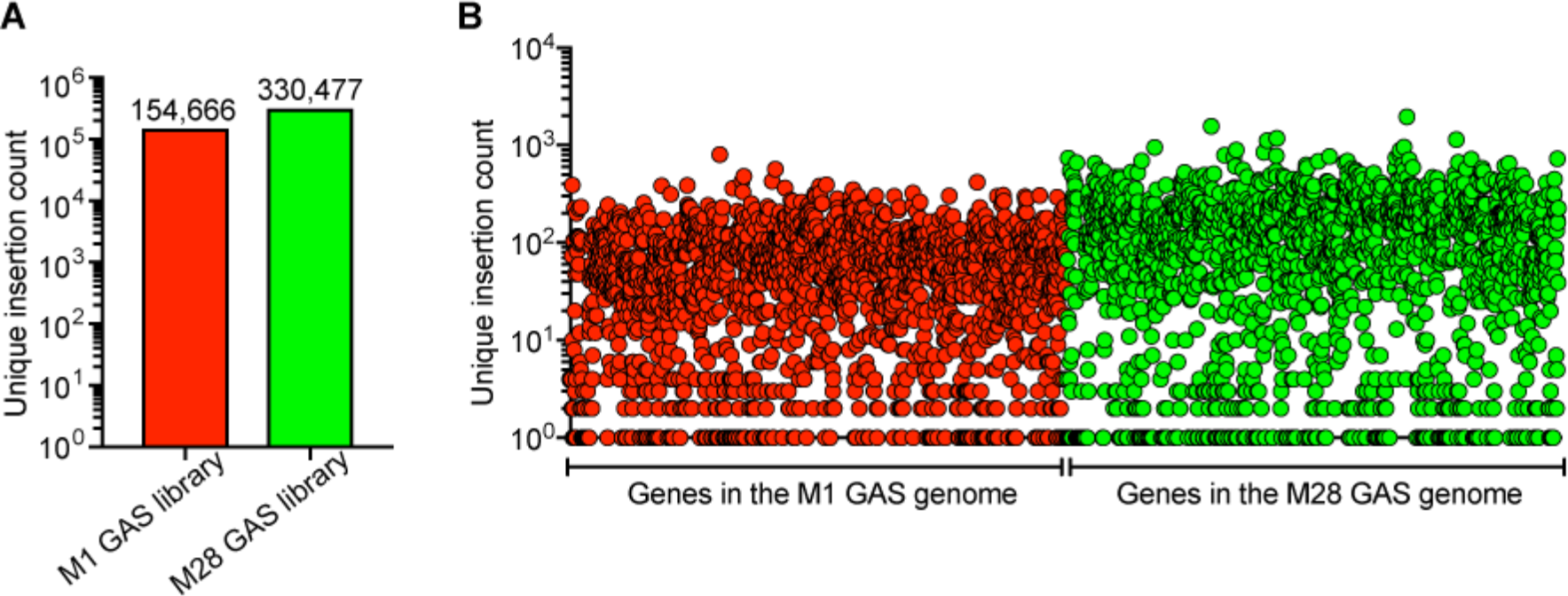
Characterization of the serotype M1 and M28 transposon mutant libraries. (**A**) Overall unique transposon insertion count of the serotype M1 (red) and M28 (green) mutant libraries. (**B**) Unique transposon insertion count of each gene in the serotype M1 (red circle) and M28 (green circle) genomes.

### Genome-wide screens identify GAS genes contributing to fitness in a necrotizing myositis infection model in cynomolgus macaques

We screened the M1 and M28 GAS transposon mutant libraries in NHPs as a first step toward discovering genes contributing to fitness during necrotizing myositis. Six cynomolgus macaques each were inoculated by intramuscular injection with either the M1 or M28 transposon mutant libraries and followed for 24 h. All animals developed signs and symptoms consistent with necrotizing myositis and were necropsied. Biopsies containing necrotic muscle were obtained from the inoculation site to recover output mutant pools for subsequent analysis. Quantitative culture yielded an average of 4.87 × 10^8^ CFU/g for M1, and 8.77 × 10^8^ for M28 in the tissue biopsy specimens used for TraDIS analysis.

TraDIS was used to compare the mutant compositions of the input and output pools. The TraDIS analysis identified genes with significantly altered mutant frequency in the output mutant pools relative to the input mutant pool (examples shown in Figure S1). Infection bottlenecks can be a technical challenge for high-throughput transposon mutagenesis studies, and substantial loss of mutant library complexity during animal infection can result in erroneous identification of fitness genes (50). Our TraDIS results showed that for both M1 and M28 GAS screens, there was no substantial decline of mutant library complexity post NHP infections (Figure 2A,B). On average, 67% and 84% of the library complexity remain in the M1 and M28 output pools, respectively (Figure 2A,B). This high diversity of transposition site mutants recovered is inconsistent with a narrow infection bottleneck and indicates that our screens are unlikely to erroneously identify fitness genes. To identify GAS fitness genes in the infected NHP skeletal muscle milieu, genes previously identified as essential for GAS growth *in vitro* in rich medium (THY) were excluded from the analysis as is commonly done (51). Disrupted genes associated with significantly decreased fitness (transposon frequency log2 fold-change < −1, and *q* value < 0.01) in the output mutant pools were interpreted as contributing to NHP necrotizing myositis (Figure 2C,D, Figure 3). We identified 126 and 116 genes in the serotype M1 and M28 strains, respectively, that are crucial for GAS fitness in this infection model (Figure 3A). That is, inactivating these genes potentially confers diminished GAS fitness in necrotizing myositis. Importantly, a common set of 72 genes was identified as crucial for fitness in both the serotype M1 and M28 library NHP screens (Figure 3A, Table S1). The shared 72 genes represent 57% of the serotype M1 fitness genes and 65% of the serotype M28 fitness genes (Figure 3A). Functional categorization of the fitness genes found that numerically, the more prevalent GOG categories included genes inferred to be involved in amino acid transport and metabolism (E), inorganic ion transport and metabolism (P), and transcription (K) (Figure 3C). Genes encoding many documented virulence factors or virulence modulating factors were identified as contributing to fitness in both serotype M1 and M28 GAS strains, including *adcB/C* (52, 53), *gacI* (54, 55), *pepO* (56, 57), *inlA* (58), *perR* (59) and *scfAB* (32) (Table S1, Table S2, Table S4).

**Figure 2.**
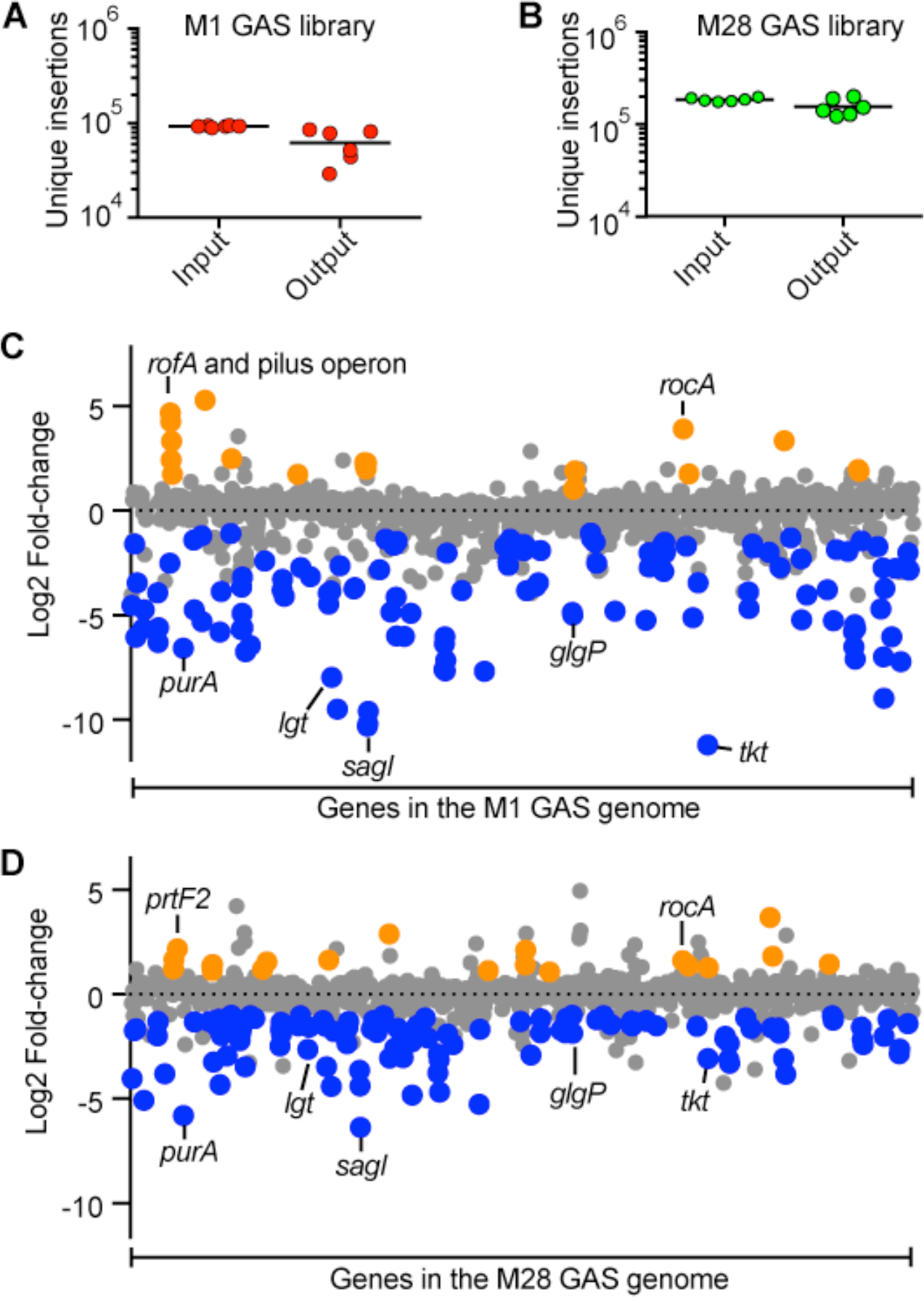
TraDIS analysis of GAS gene fitness in NHP necrotizing myositis. Complexity of the **(A)** M1 GAS mutant pools and **(B)** M28 GAS mutant pools before and after a 24-hr experimental NHP infection. Genome-scale summary of the changes in mutant abundance (*y* axis) for each of the genes (*x* axis) in the **(C)** M1 GAS output pools and **(D)** M28 GAS output pools. Gene mutations (insertions) conferring significantly decreased (blue dots) or increased (gold dots) fitness are highlighted.

**Figure 3.**
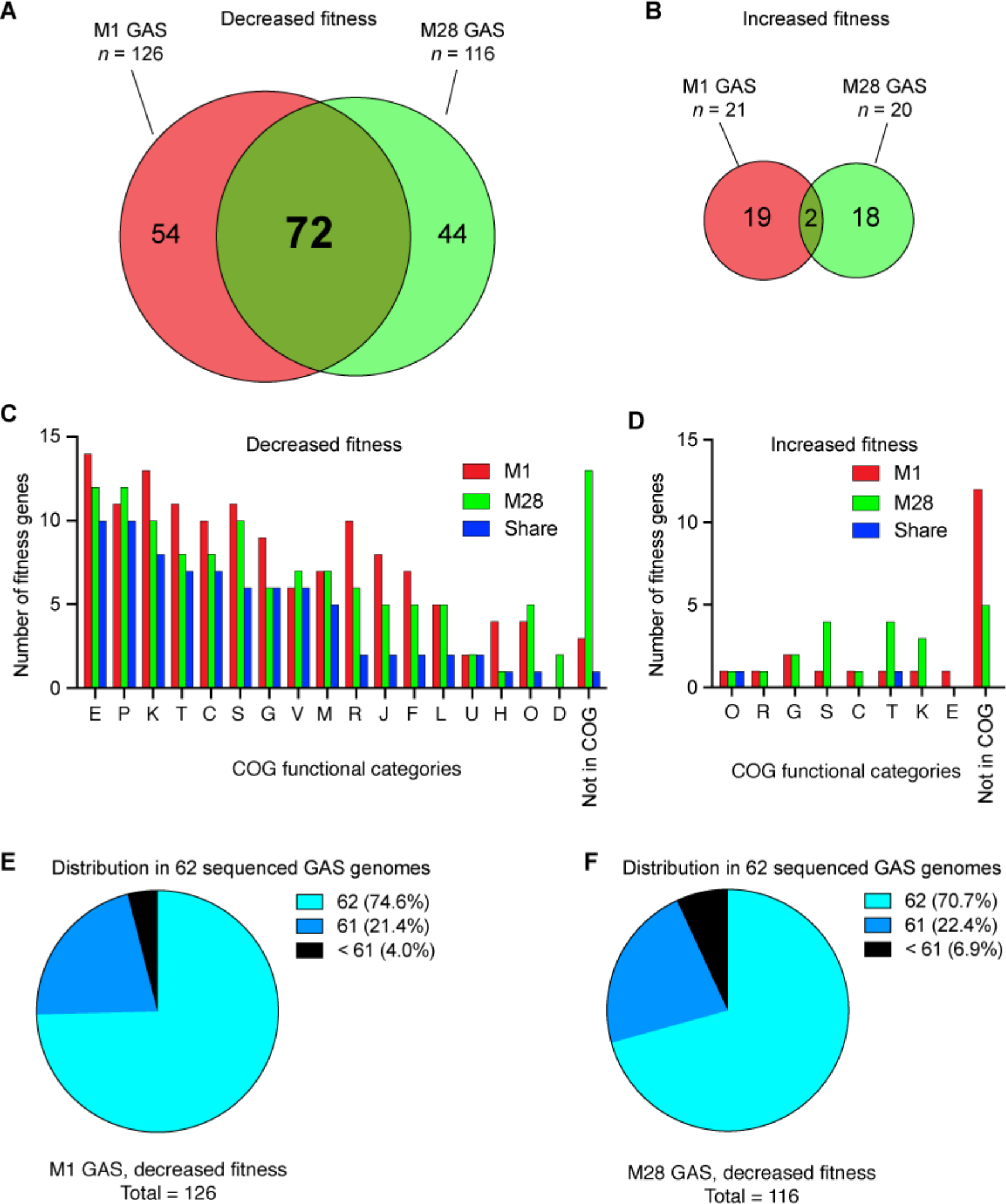
GAS gene mutations conferring significantly altered fitness during necrotizing myositis. Venn diagrams showing the number of mutated genes conferring significantly decreased fitness (**A**) and increased fitness (**B**) in M1 and M28 GAS strains during NHP infections. (**C, D**) Functional categorization of the identified GAS *in vivo* fitness genes during necrotizing myositis. (**E, F**) Distribution of the M1 and M28 GAS genes required for infection among the 62 sequenced GAS genomes. COG, clusters of orthologous groups.

We also identified 21 and 20 genes in the serotype M1 and M28 strains, respectively, that are associated with significantly increased fitness *in vivo* (transposon insertion frequency log2 fold-change > 1, and q value < 0.01) (Figure 2, Figure 3B and 3D). That is, inactivating these genes potentially confers enhanced GAS fitness during NHP necrotizing myositis. These genes include known negative regulators of virulence *rivR* (60) and *rocA* (61–65). Of note, *rocA* and *ppiB* (peptidyl-prolyl cis-trans isomerase) were identified in both the serotype M1 and M28 strains (Figure 3B, Table S3, Table S5). Inactivation of the *rocA* gene in an M28 GAS strain was recently shown to significantly increase virulence in a mouse model of necrotizing myositis (66).

To investigate the phylogenetic distribution of the identified *in vivo* fitness genes among diverse GAS strains, we examined the presence of M1 and M28 fitness genes required during NHP necrotizing myositis in 62 sequenced GAS genomes representing 26 different M protein serotypes (Table S6). The vast majority of the M1 fitness genes (96%) and M28 fitness genes (93%) are present in at least 61 of the 62 GAS strains (Figure 3, E and F).

### Comparison of the genetic requirement for necrotizing myositis and those for GAS fitness in human saliva and blood ex vivo, and mouse subcutaneous infections

Previous genome-wide transposon mutagenesis studies identified GAS genes required for growth in animal infection models and *ex vivo* in human body fluids such as blood and saliva (30–32). These published data allowed us to test the hypothesis that the GAS gene requirements for NHP necrotizing myositis are distinct from those identified by transposon screens performed in other model infection environments. That is, we were able to assess the extent to which GAS has infection-specific genetic programs. We recently reported that 92 serotype M1 genes were required for optimal growth *ex vivo* in human saliva (31). Only 19 (21% of 92) genes were defined as contributing to serotype M1 fitness in both human saliva and NHP necrotizing myositis (Figure 4A). These genes include metabolic genes (*purA*, *purB*, *acoABCL*, *glgP* and *malM*) and transporter genes (*adcAB*, *braB*, *mtsA*, *mtsB*, *artP*, and *artQ*). The great majority of M1 genes (*n* = 107, 85%) required in NHP necrotizing myositis did not overlap with fitness genes required *ex vivo* in human saliva. Using a similar transposon mutagenesis technique (Tn-seq) 147 genes were identified as contributing to fitness of serotype M1 GAS strain 5448 after subcutaneous inoculation in mice (32). The overlap between the mouse subcutaneous fitness genes and the 126 necrotizing myositis genes is relatively larger (*n* = 39) (Figure 4B). These genes include metabolic genes *purA*, *purB*, *acoABCL*, *glgP*, *malM*, *arcABCD*, and phosphotransferase system genes *manMLN*. However, 69% of the genes (*n* = 87) required for NHP necrotizing myositis did not overlap with the genes identified in the mouse subcutaneous infection study. Using a transposon mutagenesis technique, Mclver and colleagues (30) identified 81 M1 GAS genes required for optimal bacterial growth in human blood. In comparison with the 126 necrotizing myositis fitness genes, only 14 genes were required in both conditions (Figure 4C). 89% of the genes (*n* = 112) required for NHP necrotizing myositis did not overlap with the genes required in human blood *ex vivo*. These include genes for carbohydrate metabolism (*glgP*, *malM*), transporters (*adcB*, *braB*, *mtsA*, *mtsB*), and transcriptional regulators (*adcR*, *ciaH*, *ciaR*, *ihk*, *irr*). To summarize, there is only modest overlap between the GAS genes contributing to fitness during experimental NHP necrotizing myositis, relative to growth *ex vivo* in human saliva and blood, and mouse subcutaneous infection. These results are consistent with our hypothesis that GAS has infection-specific genetic programs.

**Figure 4.**
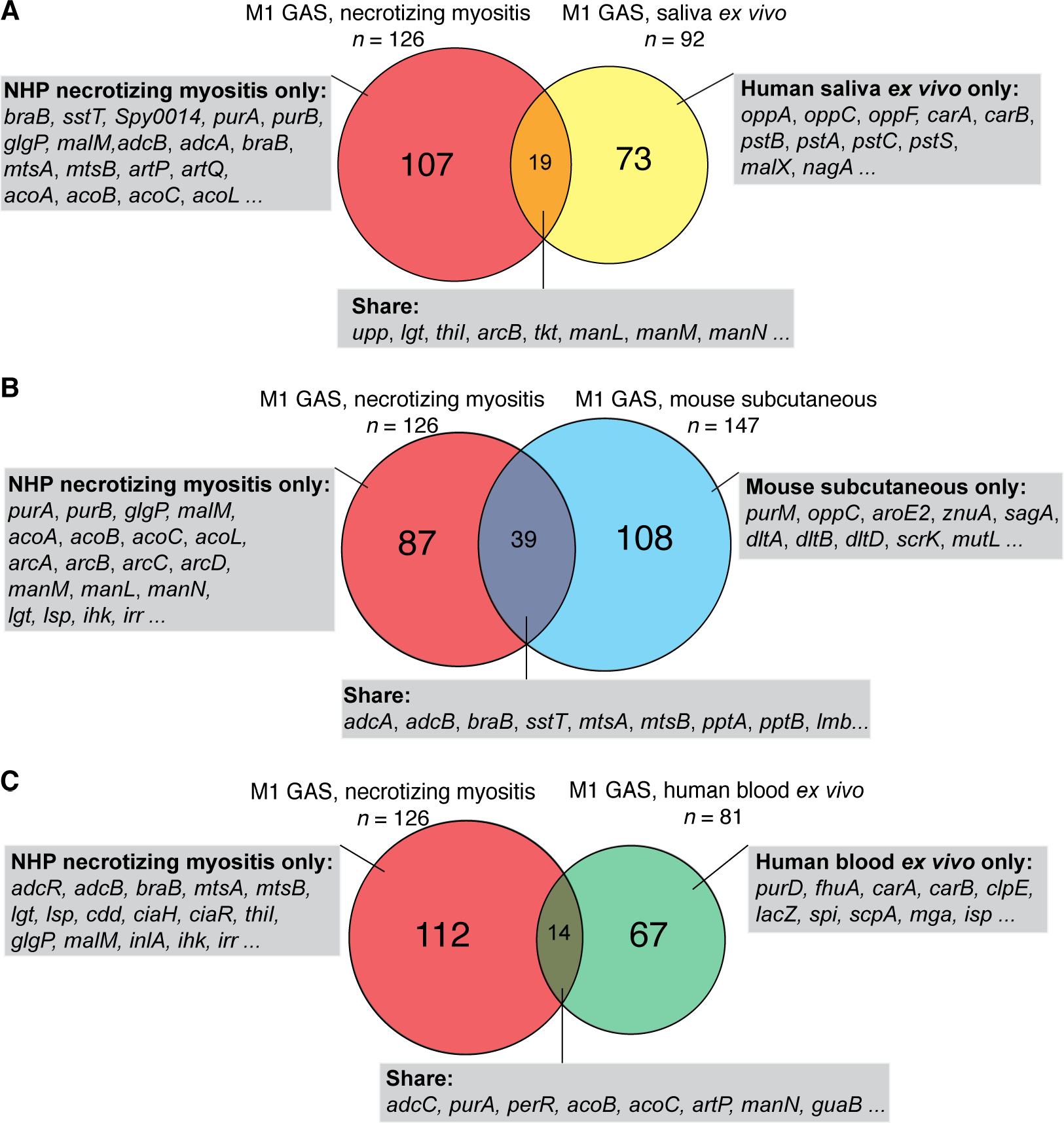
Lack of substantial overlap between GAS fitness genes required for necrotizing myositis and those required in other *in vitro* and *in vivo* environments. (**A**) Venn diagram comparison of 126 genes required for NHP necrotizing myositis with 92 genes required for optimal growth in human saliva *ex vivo*. (**B**) Venn diagram comparison of 126 genes required for necrotizing myositis with 147 genes required for mouse subcutaneous infection (32). **(C)**Venn diagram comparison of the 126 genes required for necrotizing myositis with 81 genes required for GAS growth in human blood *ex vivo*. Representative genes belonging to each category are listed in the shaded rectangular boxes (**A, B and C**).

### Genes encoding transporters constitute a considerable portion of GAS fitness determinants in experimental NHP necrotizing myositis

Bioinformatic analyses of the identified fitness genes found that regardless of M1 or M28 serotype, more than 25% of the genes contributing to *in vivo* fitness during NHP necrotizing myositis encode proven or putative transporters (Figure 5A). Specifically, 25.4 % of the serotype M1 fitness genes (*n* = 32) and 29.7% of the M28 GAS fitness genes (*n* = 32) encode proven or putative transporters (Figure 5A). Importantly, 26 transporter genes are required in both M1 and M28 strains during infection, indicating that there was considerable overlap between the two sets of genes (Figure 5B). These 26 shared genes encode 13 distinct transporters with proven or predicted roles in uptake of nutrients such as amino acids, metal ions, vitamins, carbohydrate, and export of a variety of substrates. (Figure 5C). The DNA sequences of the 26 transporter genes are highly conserved (95% to 100% identity) among genomes for 62 sequenced GAS strains representing 26 different M protein serotypes (Figure 6). One gene (*Spy0499*) has less homology in GAS (81% identical in four GAS strains) among the 62 strains with complete genomes.

**Figure 5.**
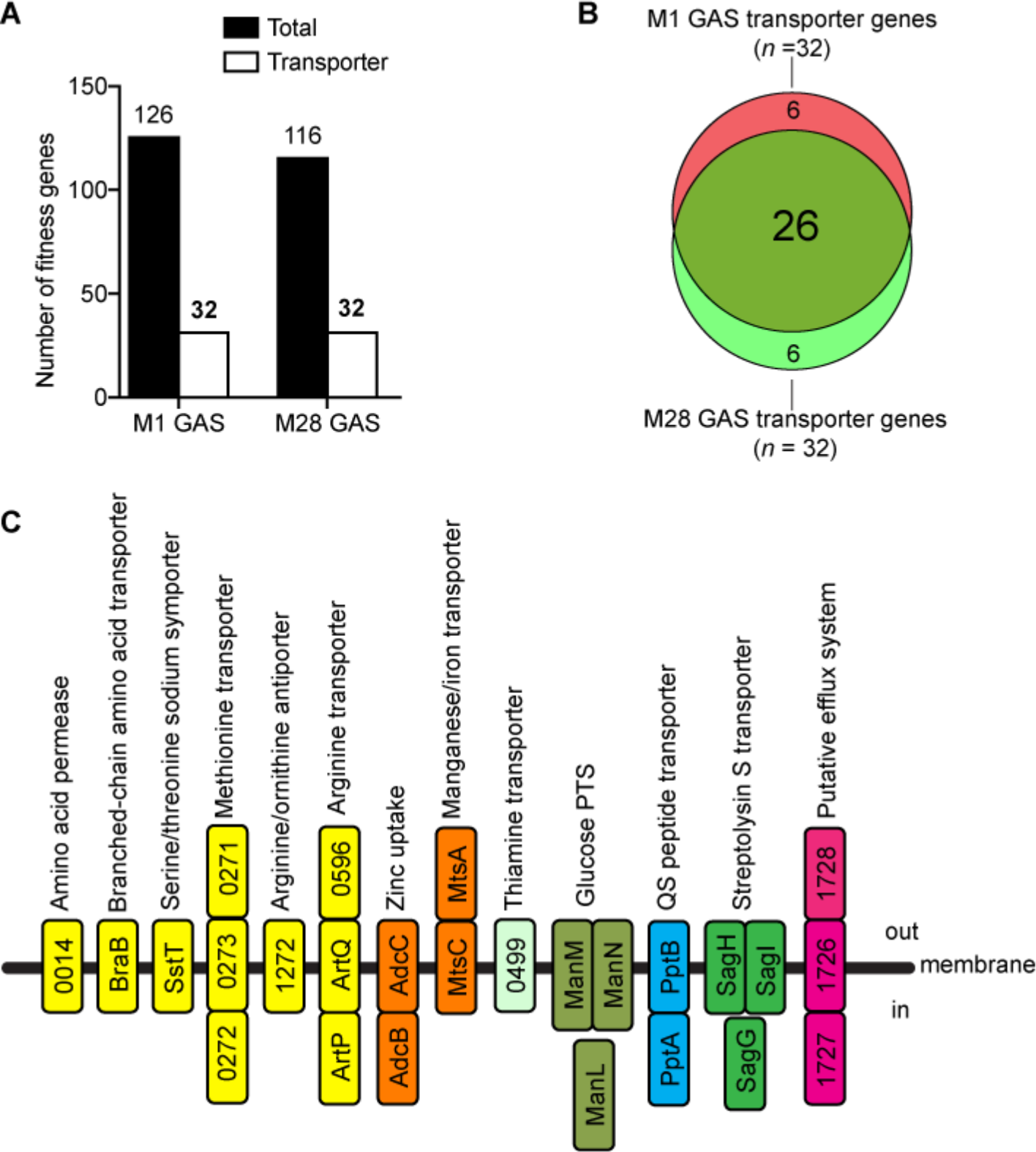
Genes encoding proven or putative transporters are an abundant portion of fitness genes that are required during necrotizing myositis in NHPs. (**A**) M1 GAS fitness genes (*n* = 32, 25.4%) and M28 GAS fitness genes (*n* = 32, 27.6%) that encode proven or putative transporters. (**B**) Venn diagram showing the relationship between M1 and M28 transporter genes required during NHP skeletal muscle infections; 26 genes are required in both M1 and M28 GAS strains. (**C**) Schematic showing the proven or putative transporters encoded by the 26 shared transporter genes and their inferred functions. Inferred transporter elements (Spy0271, Spy0596, MtsA, and Spy1728) that are likely positioned outside of the bacterial cell are putative lipoproteins. Elements that are inferred to be positioned on the membrane and in the bacterial cell are putative transmembrane proteins and cytosolic proteins, respectively. The locus tag numbers refer to the annotation for serotype M1 GAS strain MGAS5005.

**Figure 6.**
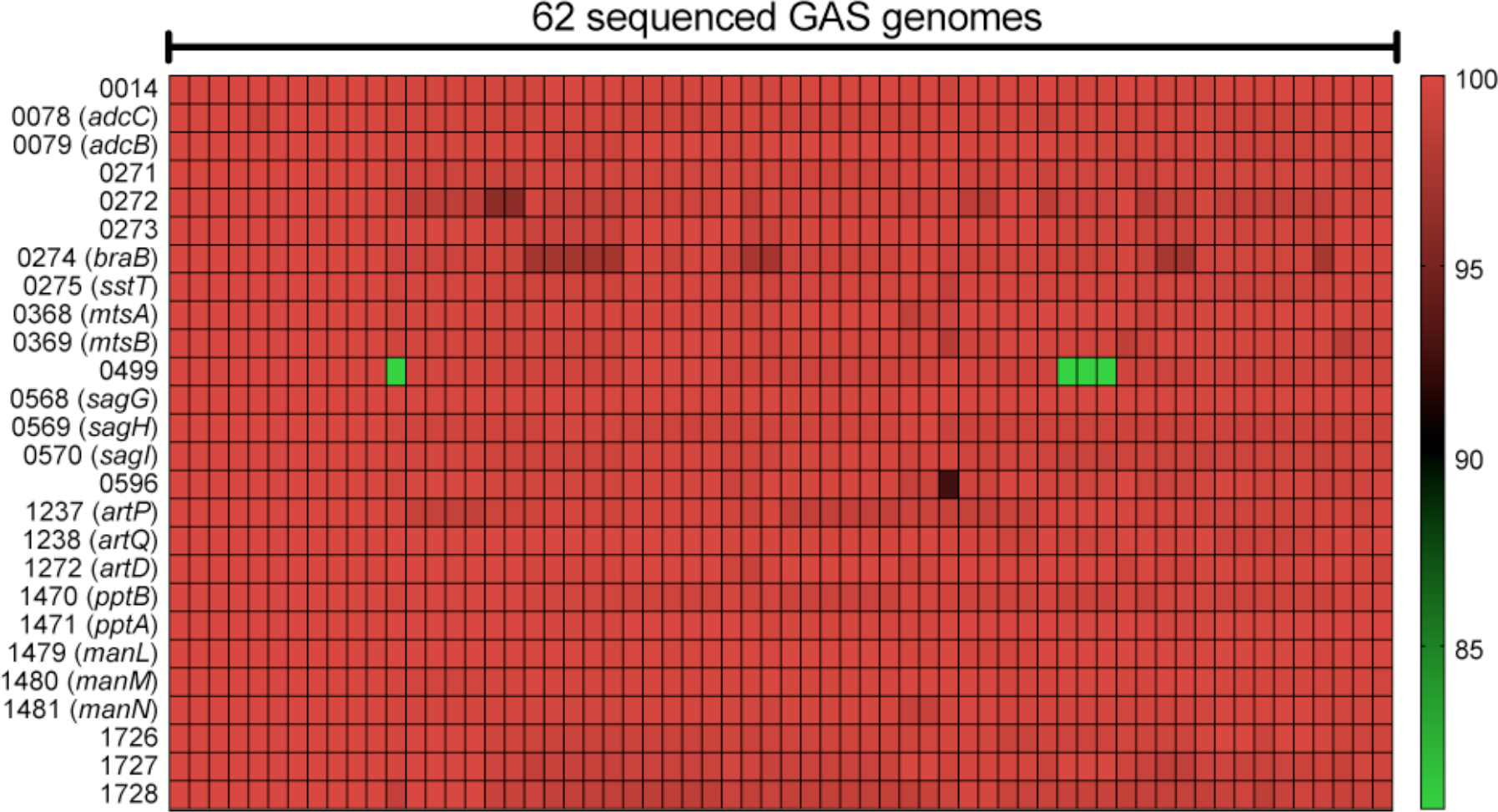
Conservation of the 26 GAS transporter genes among 62 sequenced GAS genomes. Heat map showing the percent identity of the 26 transporter genes of the 62 sequenced genomes (representing 26 different M protein serotypes) relative to those of the serotype M1 reference strain MGAS5005. The locus tag numbers refer to the annotation for serotype M1 GAS strain MGAS5005.

### Validation of the TraDIS screen results for genes encoding putative amino acid transporters

Six of the 13 transporters identified to be important for both serotype M1 and M28 strains during NHP necrotizing myositis are putative amino acid transporters (Figure 5C, yellow). For example, Spy0014 is a putative amino acid permease and BraB is a putative branched-chain amino acid transporter. Spy0271, Spy0272, and Spy0273 constitute a putative ABC transporter with similarity to methionine transporter proteins MetQ (65% identical), MetN (73% identical), and MetP (71% identical), respectively, of *Streptococcus pneumoniae* (67).

To test the hypothesis that Spy0014, BraB, and Spy0271-0273 participate in amino acid transport, we constructed isogenic deletion mutant strains Δ*Spy0014*, Δ*braB*, and Δ*Spy0271-0273* in parental M1 strain MGAS2221. We studied their growth phenotypes in rich medium (THY broth), and in a peptide-free chemically defined medium (CDM) (Figure 7). Compared to the wild-type parental strain, the three isogenic mutant strains do not have a growth defect in THY medium (Figure 7A). However, the mutant strains had a severe growth defect when cultured in CDM (Figure 7B), a result consistent with our hypothesis. As anticipated, the growth defect of these three isogenic mutant strains *ΔSpy0014*, *ΔbraB*, and *ΔSpy0271-0273* was ameliorated by supplementing CDM with 0.1 g/ml tryptone, a source of abundant peptides (Figure 7C). Together, these results are consistent with the idea that Spy0014, BraB, and Spy0271-0273 are amino acid transporters that are essential for GAS growth in the absence of a source of abundant exogenous peptides.

**Figure 7.**
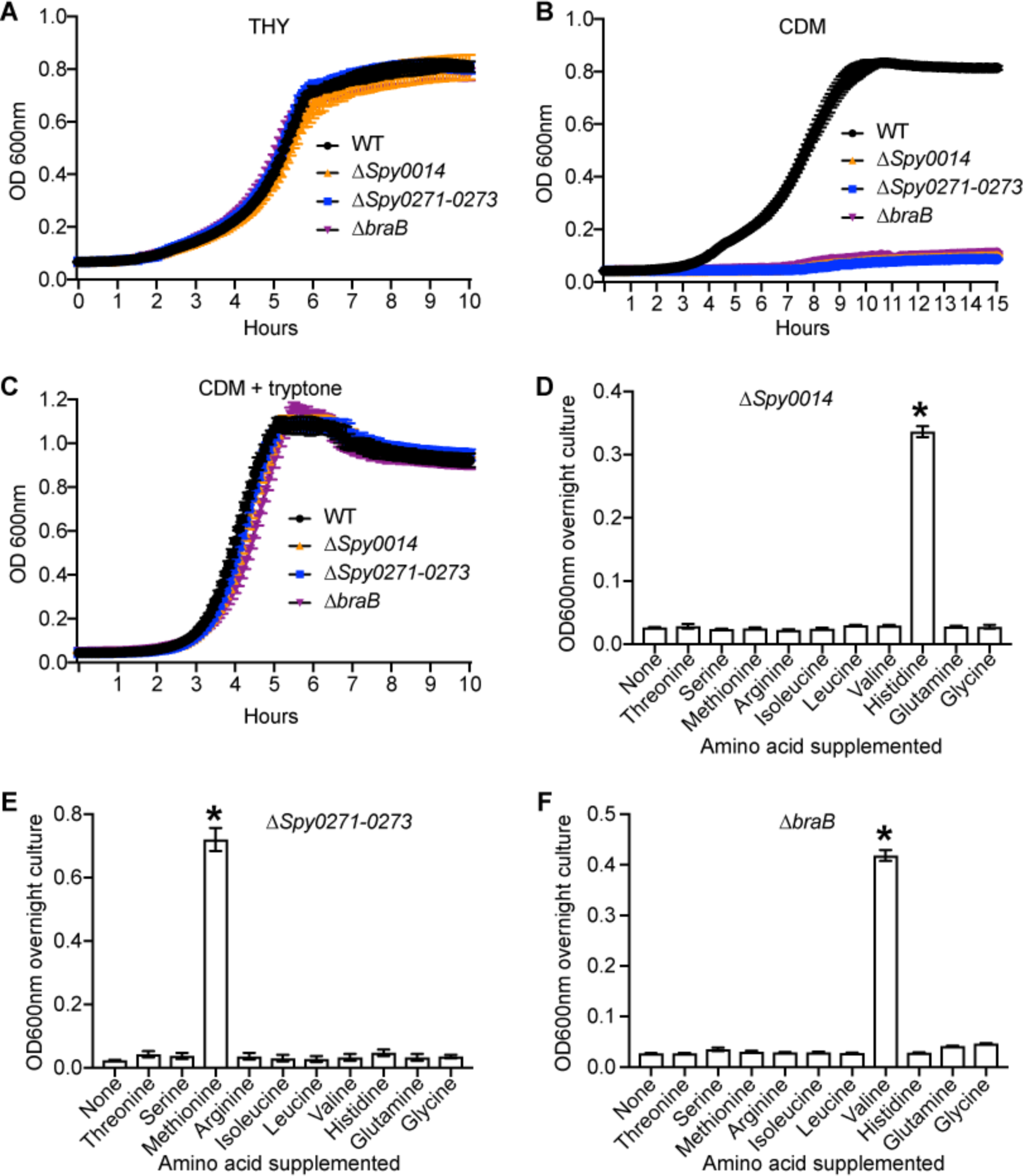
*In vitro* phenotype of three amino acid transporter mutant strains. (**A-C**) Growth of parental wild-type strain MGAS2221 (WT), Δ*Spy0014*, Δ*braB*, and Δ*Spy0271-0273* in rich medium THY (**A**), chemically defined medium (**B**), and chemically defined medium supplemented with 10g/L tryptone (**C**). (**D-F**) Growth of three mutant strains in CDM supplemented with 1g/L of specified amino acids. Experiments were performed in triplicate on 3 separate occasions. Replicate data are expressed as the mean ± SD in D, E, and F. \**P* < 0.05 vs. unsupplemented group, one-way ANOVA.

GAS is auxotrophic for 15 amino acids considered essential for growth (68, 69). We hypothesized that transporters Spy0014, BraB and Spy0271-0273 contribute to uptake of specific essential amino acids. To test this hypothesis, we supplemented CDM with a high concentration (1 g/L) of each of the highly soluble essential amino acids to determine if certain amino acids restored the growth of these transporter mutants via non-specific uptake (Figure 7D-F). Consistent with the hypothesis, supplementing CDM with methionine restored the growth of mutant *ΔSpy0271-0273* to near-wild-type growth phenotype, suggesting that *Spy0271-0273* encode a methionine transporter (Figure 7E). Similarly, supplementing CDM with histidine and valine partially restored the growth of mutant strains *ΔSpy0014* and *ΔbraB*, respectively, suggesting that Spy0014 and BraB contribute to uptake of histidine and valine (Figure 7, D and F).

### Validation of the TraDIS screen results in the NHP model of necrotizing myositis

To validate the TraDIS screen results *in vivo*, we infected NHPs in the quadriceps with parental M1 strain MGAS2221, and isogenic mutant strains Δ*Spy0014*, Δ*braB*, and Δ*Spy0271-0273*. Compared to the wild-type parental strain, each of these three transporter mutant strains caused significantly smaller lesions characterized by less tissue destruction in the NHP necrotizing myositis model (Figure 8, A and B). In addition, compared to the wild-type parental strain, significantly fewer CFUs of each isogenic mutant strain were recovered from the inoculation site and a distal site of dissemination (Figure 8, C and D).

**Figure 8.**
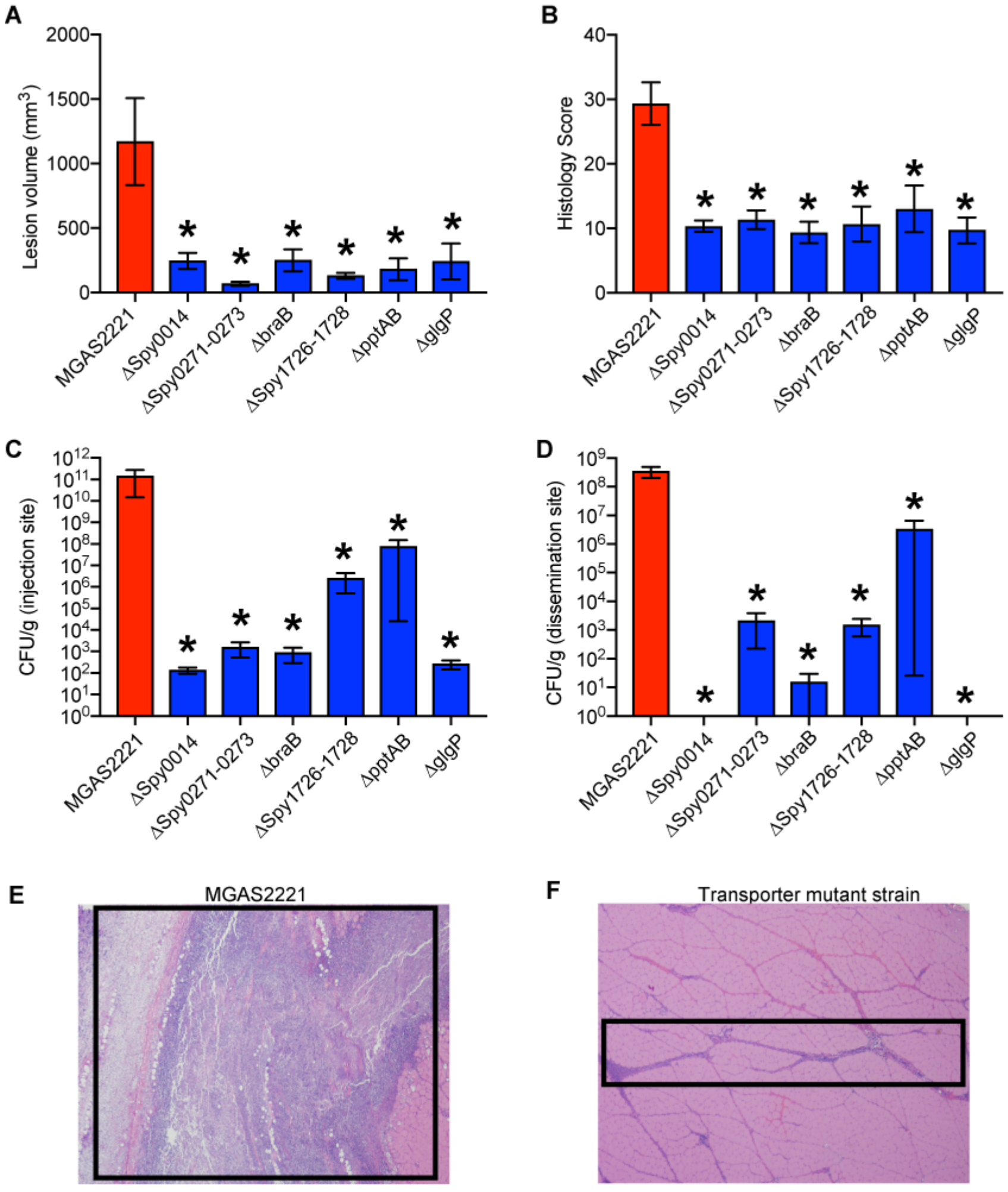
Virulence phenotypes of GAS isogenic transporter deletion mutant strains in NHPs. (**A,B**) Volume (left) and histology score (right) of the necrotizing myositis lesions caused by the parental wild-type M1 GAS strain MGAS2221 compared to each isogenic deletion mutant strain. (**C,D**) Colony forming units recovered from the inoculation site (left) and distal muscle margin (right). For all panels, mean ± SEM is shown. \**P*<0.05, Mann-Whitney test (panels A, C and D) or Wilcoxon Rank Sum test (panel B). Micrographs of hematoxylin and eosin necrotizing myositis lesions caused by the parental wild-type stain (E) compared to a representative transporter mutant strain Δ*Spy0014* (F). The boxes enclose each necrotic lesion (original magnification 2x).

### Virulence role of Spy1726-1728, a poorly characterized ABC transporter

Our genome-wide screens identified a putative ABC transporter of unknown function that is required for NHP infection by the M1 and M28 GAS strains (Figure 5C, red). This putative transporter is encoded by three contiguous genes: Spy1726 (transporter permease protein), Spy1727 (ATP-binding protein), and Spy1728 (substrate-binding lipoprotein). To confirm the virulence role of this putative transporter, we used isogenic mutant strain Δ*Spy1726-1728* made by deleting the entire *Spy1726-1728* region in serotype M1 parental strain MGAS2221. Consistent with the result from the initial NHP necrotizing myositis TraDIS screen, isogenic mutant strain Δ*Spy1726-1728* is significantly attenuated in capacity to cause necrotizing myositis in NHPs (Figure 8, A - D). This putative ABC transporter operon was not identified as important for virulence in previous GAS transposon mutagenesis screens (24, 30, 32), suggesting an infection- or primate-specific role in necrotizing myositis. Of note, Spy1728 (a substrate-binding lipoprotein) was previously shown to be a GAS surface protein and potential vaccine candidate (70).

### Virulence role of quorum sensing peptide transporter PptAB

Our genome-wide TraDIS screens suggest that inactivating the quorum sensing peptide transporter PptAB in the serotype M1 and M28 strains results in significantly decreased fitness in NHP necrotizing myositis. To test this finding, we generated isogenic mutant strain *ΔpptAB* by deleting the *pptAB* genes in serotype M1 parental strain MGAS2221. Relative to the WT parental strain, the *ΔpptAB* mutant strain is significantly attenuated in ability to cause necrotizing myositis in NHPs (Figure 8, A-D). PptAB has been reported to be required for exporting the SHP2 and SHP3 quorum sensing peptides (71). However, Rgg2, the transcriptional regulator that controls expression of the *shp2* and *shp3* genes was not identified as important for NHP infections in our TraDIS screens. These results suggest the attenuation of the virulence phenotype in the Δ*pptAB* mutant strain is likely not associated with the SHP2, SHP3 quorum-sensing pathway in serotype M1 and M28 strains in this infection model.

### *Confirmation of the virulence role of* glgP, *a gene involved in carbohydrate utilization*

Our TraDIS screens identified many GAS genes implicated in transport of nutrients such as amino acids, vitamins and carbohydrates. We next studied a gene likely to be involved in carbohydrate utilization. The GAS gene *glgP* was identified as essential for necrotizing myositis in the TraDIS screens conducted with both the serotype M1 and M28 transposon mutant libraries (Table S1). However, *glgP* has not previously been implicated in GAS virulence. *glgP* encodes an inferred protein with homology to *E. coli* glycogen phosphorylase (72). We generated isogenic mutant strain Δ*glgP* by deleting the *glgP* gene in serotype M1 strain MGAS2221. The Δ*glgP* isogenic mutant strain is severely attenuated in capacity to cause necrotizing myositis in NHPs, thereby confirming the TraDIS screen finding (Figure 8, A-D). We next evaluated the potential role of *glgP* in carbohydrate metabolism. Although the Δ*glgP* mutant strain has no growth defect in medium with glucose, this mutant strain has a severe growth defect when maltose or maltodextrin is provided as the sole carbohydrate in the culture medium (Figure 9, A-C). Consistent with the idea that the product of the *glgP* gene is involved in glycogen and starch utilization, bacteria grown in THY supplemented with starch showed evidence of starch accumulation in the isogenic mutant strain Δ*glgP*, but not the wild-type parental strain that retains the ability to metabolize starch (Figure 9D). Interestingly, in *E. coli*, glycogen accumulation is also significantly higher in *glgP* deletion mutants (72).

**Figure 9.**
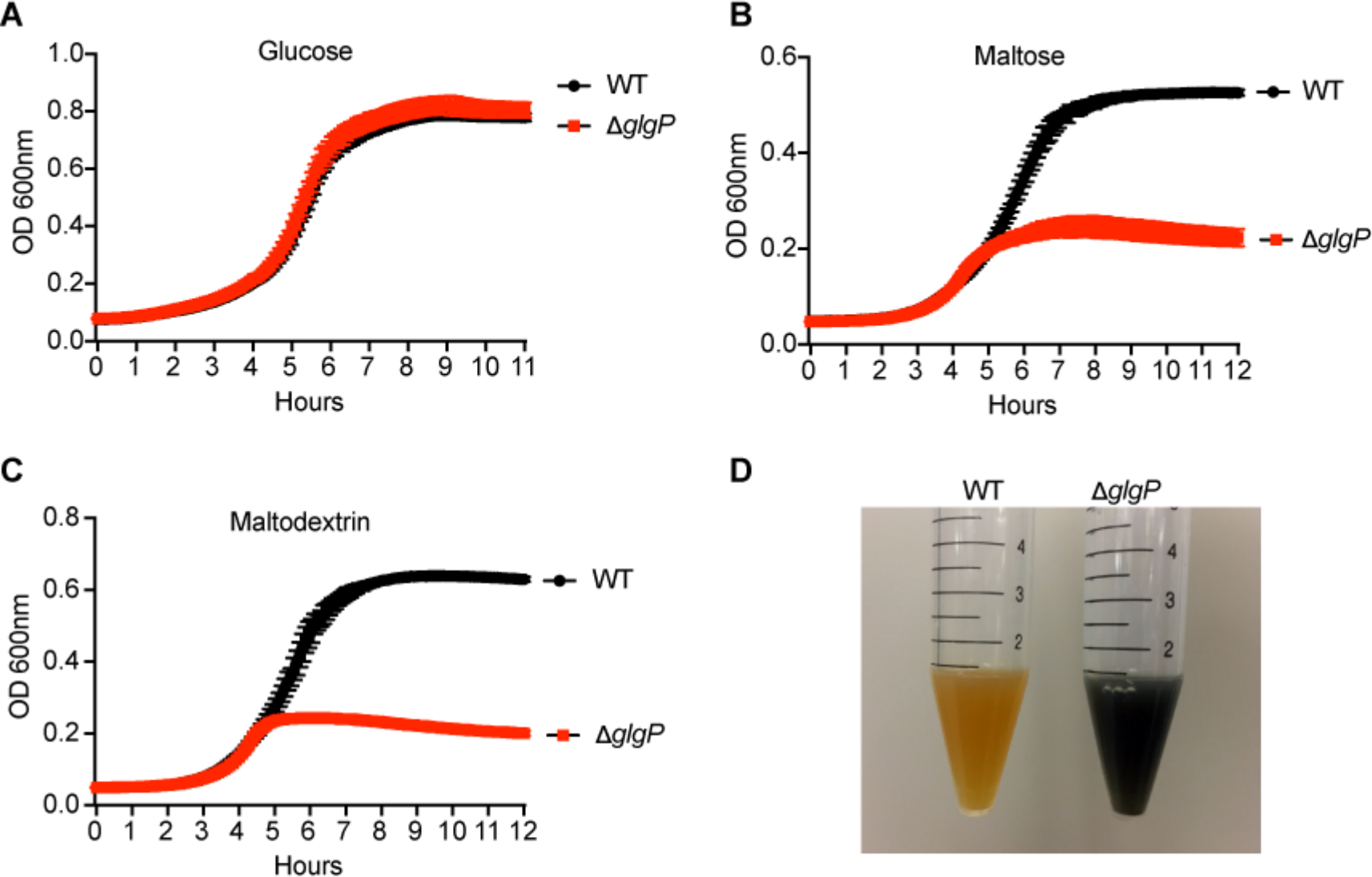
*In vitro* phenotype of isogenic mutant strain Δ*glgP*. Growth of WT and isogenic mutant Δ*glgP* in THY broth with glucose as the sole carbohydrate source (**A**), THY with maltose as the sole carbohydrate source (**B**), and THY with maltodextrin as the sole carbohydrate source (**C**). (**D**) Accumulation of starch by the Δ*glgP* mutant strain.

### Expression of the GAS genes implicated in in vivo fitness genes during NHP necrotizing myositis

In the aggregate, data from the *in vivo* transposon mutant library screens and analysis of the isogenic mutant strains imply that the genes identified are expressed during NHP necrotizing myositis. To directly test for expression *in vivo*, we used TaqMan qRT-PCR to measure the transcript level of GAS transporter genes *Spy0014*, *Spy0271*, *braB*, *Spy1726*, *pptA*, and metabolic gene *glgP* in the NHP muscle tissue infected with M1 GAS MGAS2221. The transcript of all six of the GAS fitness genes studied were detectable by TaqMan qRT-PCR, thereby confirming that these genes are expressed *in vivo* in NHP necrotizing myositis (Figure 10A).

**Figure 10.**
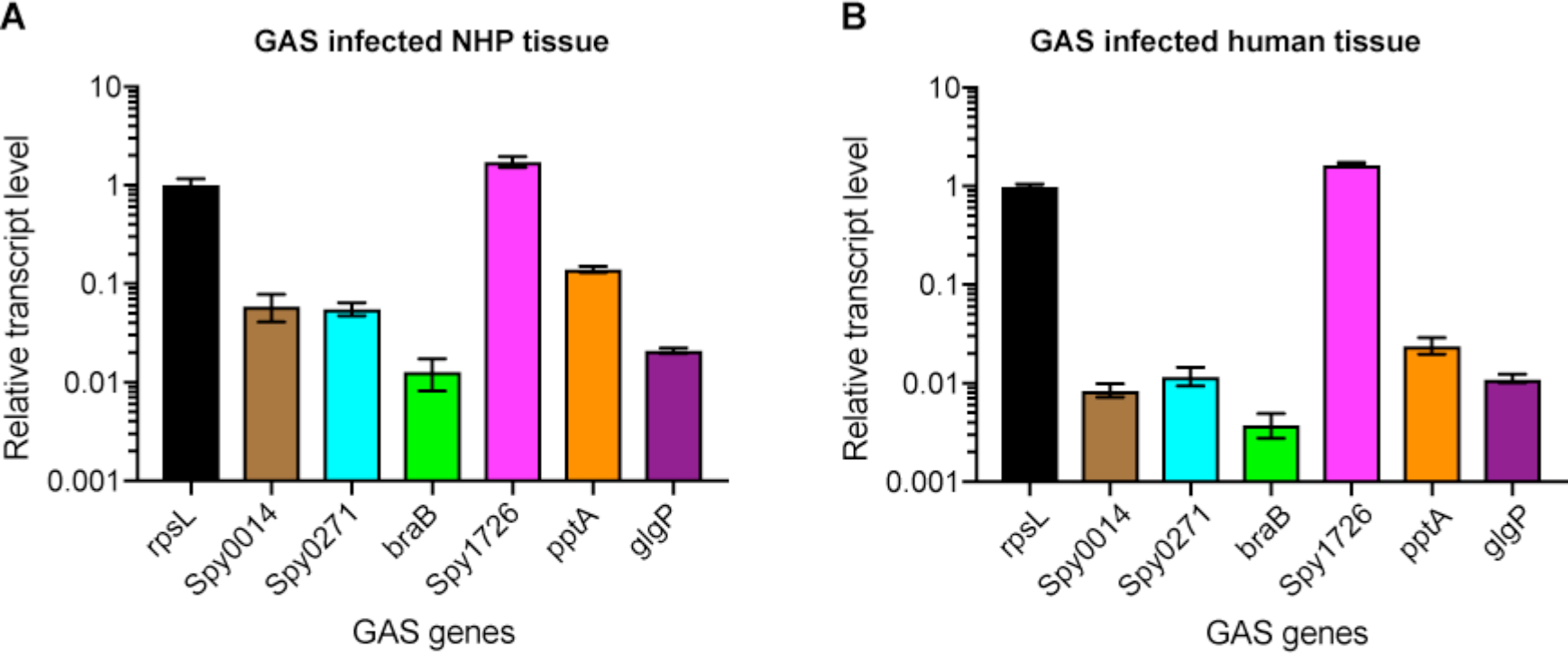
Relative transcript level of GAS fitness genes in NHP necrotizing myositis (A) and in an infected human with necrotizing fasciitis (B). *In vivo* transcript level of GAS genes *Spy0014*, *Spy0271*, *braB*, *Spy1726*, *pptA*, and *glgP* relative to housekeeping gene *rpsL*. The experiment was performed in triplicate, and mean ± SD are shown.

### Expression of fitness genes in vivo in a human with necrotizing myositis

We next tested the hypothesis that the six targeted genes of interest are expressed in a human patient with necrotizing myositis. Necrotic skeletal muscle obtained from a patient with culture-proven GAS infection was studied by TaqMan qRT-PCR. The results confirmed the presence of transcripts from the six genes in the infected human patient (Figure 10B). Important to note, the relative transcript levels for all genes tested were closely similar in the experimentally infected NHPs and humans with natural infection.

## Discussion

GAS is an abundant human pathogen that is responsible for substantial human illness and economic loss worldwide. Necrotizing fasciitis and myositis caused by this organism are particularly devastating infections because they have high morbidity and mortality. Effective treatment options for these infections remain limited and a licensed human GAS vaccine is not available. Thus, a fuller understanding of pathogen factors that contribute to these severe diseases is warranted and may facilitate translational research activities.

Our genome-wide screens identified 126 M1 genes and 116 M28 genes contributing to fitness in NHP necrotizing myositis. Of particular importance, we discovered a significant overlap between the genes identified in the M1 and M28 *in vivo* fitness screens, with 72 genes common to both serotypes, representing 57% and 64% of the M1 and M28 fitness genes, respectively. The similarity between M1 and M28 *in vivo* fitness gene results suggests the existence of conserved programs used by multiple diverse GAS strains to proliferate and damage tissue in necrotizing myositis. Many of the shared 72 genes encode proven or putative metabolic enzymes implicated in complex carbohydrate metabolism (*malM* and *glgP*), pyruvate metabolism (*acoA*, *acoB*, *acoL*), amino acid biosynthesis (*tkt*, *aroD*, *glnA*, and *arcB*), and nucleotide biosynthesis (*purA* and *purB*), suggesting these pathways are critical for GAS fitness in the environment of deep-tissue infection. In addition to metabolic genes, several previously identified GAS virulence or fitness factors were also among the 72 genes. For example, ScfAB (two putative membrane proteins) were identified as important for GAS fitness and virulence during subcutaneous infection in mice (32). *adcABC* (zinc importer) is critical for GAS virulence in mice and has vaccine interest (52).

In contrast to the similar critical gene requirements for serotype M1 and M28 GAS during experimental NHP necrotizing myositis, there is relatively little overlap between M1 GAS genes required for necrotizing myositis and growth in human saliva *ex vivo* (Figure 4). That is, the spectrum of genes contributing to fitness in these two environments is largely distinct. For example, genes encoding multiple amino acid transporters (*e.g.*, *Spy0014*, *braB* and *sstT*) required for GAS fitness during necrotizing myositis were not identified as important for growth *ex vivo* in human saliva (Figure 4A). In contrast, a GAS phosphate transporter encoded by the *pst* operon is essential for persistence in human saliva *ex vivo* but is apparently dispensable for NHP muscle infection (Figure 4A). These results suggest that amino acid uptake is critical for GAS fitness during muscle infections, whereas phosphate uptake is essential for growth in human saliva *ex vivo*. Similarly, GAS genes contributing to NHP necrotizing myositis and mouse subcutaneous infection also have relatively little overlap (Figure 4B). Many GAS metabolic genes are specifically required for NHP necrotizing myositis. For example, genes for *de novo* purine nucleotide biosynthesis (*purA* and *purB*), carbohydrate utilization (*glgP* and *malM*), and arginine and citrulline catabolism (*arcABCD*) are uniquely important for NHP necrotizing myositis. Moreover, several known streptococcal virulence-modulating factors were also identified in NHP necrotizing myositis. These include genes for GAS lipoprotein processing (*lgt* and *lsp*) (73–75), and genes for a two-component regulatory system that is essential for GAS to evade human innate immunity (*ihk* and *irr*) (76, 77) (Figure 4B). Taken together, these results suggest the nutritional environment and the survival pressures present in the infected NHP skeletal muscle are distinct from those in human saliva *ex vivo* and in a mouse subcutaneous infection model. These results imply that complex gene programs used by GAS to cause other types of human infections (e.g., puerperal sepsis and pharyngitis) are also likely to be niche-specific.

A key theme of our M1 and M28 NHP genome-wide screens was identification of many genes encoding transporters that are required during necrotizing myositis. Pinpointing exactly which of the transporters contribute to bacterial virulence during necrotizing myositis sheds new light on the mechanisms of GAS-host interactions in this severe infection. We identified 13 distinct transporters that are required in both M1 and M28 GAS strains. Six of these transporters are inferred to function in amino acid transport. This observation suggests the ability to efficiently acquire host amino acids is critical for the pathogenesis of GAS necrotizing myositis. *In vitro* growth assays showed that amino acid transporter mutant strains Δ*Spy0014*, Δ*braB*, and Δ*Spy0271-0273* have a significant growth defect in the peptide-free CDM that is ameliorated by supplementing CDM with tryptone, a source of abundant peptides. These results suggest that efficient amino acid uptake is critical for wild-type GAS growth when the peptide source is limited, and the nutritional environment of the infected muscle is probably a poor source of available peptides. GAS is auxotrophic for 15 amino acids considered essential for growth (68). We hypothesized that transporters Spy0014, BraB and Spy0271-0273 are required for the highly efficient uptake of certain essential amino acids. As anticipated, we showed that supplementing CDM with methionine, histidine, and valine restored the growth of mutant strains Δ*Spy0271-0273*, Δ*Spy0014* and Δ*braB*, respectively, indicating these three transporters contribute to transport of these three amino acids. Interestingly, the concentration of free methionine, histidine, and valine in the human skeletal muscle tissue are 16.4 mg/L, 57.4 mg/L, and 30.5 mg/L, values lower than those present in CDM (100 mg/L) (78, 79). In the aggregate, our results suggest efficient uptake of essential host amino acids such as methionine, histidine, and valine is important for GAS to cause necrotizing myositis in NHPs. The qRT-PCR data demonstrating the presence of transcripts from these transporter genes document that they are expressed in NHPs and infected humans. Together, this implies that blocking the uptake of essential amino acids by GAS might be a feasible strategy to control GAS infection pathology, but further studies are required to test this idea.

We identified several virulence-related transporters with unclear functions. One example is the putative ABC transporter comprised of Spy1726 (transporter permease protein), Spy1727 (ATP-binding protein), and Spy1728 (substrate-binding lipoprotein). The TraDIS screen results and the virulence phenotype of the isogenic mutant strain indicate this ABC transporter of unknown function is critical for the ability of GAS to cause NHP skeletal muscle pathology. Although its function is not known, multiple additional leads indicate ABC transporter Spy1726-1728 plays a role in host-pathogen interactions. For example, the Spy1726-1728 operon is upregulated when GAS cells are in human blood and macrophages (80, 81). In addition, Spy1726-1728 is regulated by the CovR/S two-component system, a global virulence gene regulator in GAS (82). Proteomic studies show that the substrate–binding lipoprotein Spy1726 is located on the bacterial cell surface, and thus might be a candidate for vaccine or other translational research (70). Future structural and functional studies on this transporter appear to be warranted.

To summarize, in this work we used two distinct transposon mutant libraries made in serotypes of GAS that cause abundant human cases of severe invasive infections to identify genes contributing to GAS fitness in an NHP model of necrotizing myositis. NHP infection using six isogenic mutant strains confirmed the crucial requirement for the genes identified by our TraDIS screens. Our findings complement work conducted with other transposon mutant screens that identified GAS genes contributing to fitness during growth *in vitro*, in human saliva and human blood *ex vivo*, and mouse subcutaneous infection. The findings presented herein may ultimately lead to better ways to diagnose, treat, and prevent necrotizing myositis and fasciitis caused by GAS, infections with devastating consequences to the human host.

## Methods

### Bacterial strains

Strain MGAS2221 is genetically representative of the pandemic clone of serotype M1 GAS that arose in the 1980s and has spread worldwide (20). Strain MGAS27961 is genetically representative of a virulent serotype M28 clone that is prevalent in the United States and elsewhere (44). These two strains have wild-type alleles of all major transcriptional regulators known to affect virulence, such as *covR* and *covS*, *ropB*, *mga*, and *rocA*.

### GAS strain growth conditions

The GAS strains were cultured in Todd-Hewitt broth supplemented with 0.5% yeast extract (THY). When required, GAS strains were grown in a chemically defined medium (CDM) (79). Amino acids were added to the designated concentration. For growth in medium with a carbohydrate source other than glucose, GAS strains were cultured in maltose medium and maltodextrin medium. The composition of these media is presented in Table S8.

### Transposon mutant libraries and culture conditions

The mutant library generated in serotype M1 strain MGAS2221 using transposon plasmid pGh9:IS*S1* was recently described (31, 49). The serotype M28 strain MGAS27961 transposon mutant library was made by essentially identical methods. The strains were grown in Todd-Hewitt broth supplemented with 0.2% yeast extract (THY) broth at 37° C with 5% CO_2_.

### Preparation of transposon mutant library frozen stock for nonhuman primate infection

100 µl of the stock transposon mutant library (M1 or M28 GAS library) was inoculated in 500 ml THY supplemented with 0.5 µg/ml erythromycin and cultured at 37° C for 8 hrs. The proliferated transposon library was pelleted by centrifugation, washed three times with saline, and then suspended in 10 ml saline supplemented with 20% glycerol. The suspended mutant library was aliquoted into cryogenic tubes and stored at −80° C until subsequent use in NHP infections.

### Nonhuman primate necrotizing myositis infection model used for TraDIS analysis

A well-described NHP model of necrotizing myositis was used (18, 33). For the transposon mutant library screens, six cynomolgus macaques (1-3 years, 2-4 kg, males and females) each were used for the serotype M1 and M28 screens. Briefly, NHPs were sedated and bacteria were inoculated in the right quadriceps. Animals were observed and necropsied 24 hours post-infection. To analyze the output transposon insertion library, a ~0.5 g (~0.5-1.0 cm diameter) biopsy of necrotic muscle was obtained from the inoculation site, homogenized in 1 ml sterile PBS, transferred to 40 ml Todd Hewitt broth and incubated for 6 hrs. Before incubation, 100 µl were removed from the 40 ml culture, serially diluted in sterile PBS and plated to determine the number of CFUs in the output library.

Infected tissue was also collected for histologic examination. The inoculum used for the input M1 transposon mutant library was ~5×10^8^ CFU/kg. The inoculum used for the input M28 transposon mutant library was 1×10^10^ CFU/kg. A higher dose of the M28 input transposon mutant library was used because in previous NHP studies, an approximately 10-to-100-fold higher inoculum of M28 strains compared to M1 strains was needed to generate similar disease character.

### DNA preparation and massively parallel sequencing

The mutant library genomic DNA preparation and DNA sequencing were performed according to procedures described previously for TraDIS analysis (31, 49). The PCR-amplified libraries were sequenced with a NextSeq550 instrument (Illumina) using a single-end 75-cycle protocol.

### Processing of TraDIS sequencing reads and data analysis

The processing of TraDIS reads and data analysis were performed according to previously described procedures (31). Briefly, the multiplexed raw Illumina reads obtained from the input and output mutant pools were parsed with FASTX Barcode Splitter (http://hannonlab.cshl.edu/fastx_toolkit/commandline.html). The resulting sequencing reads were analyzed with the TraDIS toolkit (83). tradis_comparison.R was used to compare the reads mapped per gene between the input pools (mutant libraries before NHP infection) and the output pools (mutant libraries recovered from infected NHP). The GAS genes with significantly changed mutant frequency (log2 fold-change greater than +/−1, and *q* value < 0.01) in the output mutant pools were interpreted as contributing to GAS fitness during NHP necrotizing myositis. Illumina sequencing reads of the M1 input library (*n* = 6), M1 output library (*n* = 6), M28 input library (*n* = 6), and M28 output library (*n* = 6) are deposited in the NCBI Sequence Read Archive (SRA) under the accession number (xxxxxxxxxx).

### Construction and characterization of isogenic mutant strains

Isogenic mutant strains were derived from wild-type parental strain MGAS2221, the organism used for construction of the serotype M1 transposon mutant library. Primers used for generating the mutant strains are listed in Table S7. Markerless isogenic mutant strains were constructed by nonpolar deletion of the target gene(s) using allelic exchange (18). For example, to delete *Spy0014*, primer sets 0014-1/2 and 0014-3/4 were used to amplify two ~1.5 kb fragments flanking *Spy0014* with genomic DNA purified from serotype M1 strain MGAS2221. The two flanking fragments were combined by overlap-extension PCR with primers 0014-1 and 0014-4. The combined fragment was cloned into suicide vector pBBL740 and transformed into parental strain MGAS2221. The plasmid integrant was used for allelic exchange as described previously (18). PCR was used to identify potential mutant candidates containing the desired deletion. All other isogenic mutant strains were generated using analogous methods. Whole genome sequencing of all isogenic mutant strains was done to confirm the absence of spurious mutations.

### Infection of NHPs with isogenic mutant strains

To confirm the role of candidate genes in necrotizing myositis molecular pathogenesis and thereby validate the TraDIS data, the virulence of the parental wild-type strain MGAS2221 and the six isogenic deletion-mutant strains was assessed in the NHP necrotizing myositis infection model. Animals randomly assigned to different strain treatment groups received 10^8^ CFU/kg of one strain (wild-type or isogenic mutant) in the right limb and a different strain in the left limb. Each strain was tested in triplicate. The animals were observed continuously and necropsied at 24 hrs post-inoculation.

### Histopathology analysis

For histology evaluation, lesions were excised, and visually inspected. Lesions (necrotic muscle) were measured in three dimensions and volume was calculated using the formula for an ellipsoid. Tissue taken from the inoculation site was trisected, fixed in 10% phosphate buffered formalin, and embedded in paraffin using standard automated instruments. Histology of the three sections taken from each limb was scored by a pathologist blinded to the strain treatment groups as described previously (18, 19). To obtain the quantitative CFU data, diseased muscle obtained from the inoculation site or distal hip margin was weighed, homogenized (Omni International) in 1 mL PBS, and CFUs were determined by plating serial dilutions of the homogenate. Statistical differences between strain groups were determined using the Mann-Whitney test.

### *Iodine staining of wild-type and* ΔglgP *mutant strain*

The wild-type strain and isogenic Δ*glgP* mutant strain were cultured for eight hours in 10 ml of THY supplemented with 2g/L of soluble starch (Sigma-Aldrich). GAS cells were pelleted and washed five times with saline to remove the culture medium. After the final saline wash, GAS cells were suspended in 1ml of saline. 10 µl of Gram’s iodine solution was added to the cell suspensions to visualize glycogen accumulation in the GAS cell. Only the Δ*glgP* mutant strain displayed a dark blue iodine stain phenotype.

### Isolation of total RNA from GAS-infected non-human primates quadriceps muscle sections and skeletal muscle from a human with GAS necrotizing fasciitis

Infected tissue from NHPs or human patients was stored at −80°C in DNA/RNA Shield (Zymo Research) or RNAlater (Invitrogen), respectively, thawed on ice, transferred to a tube containing 2 ml of cold TE, and diluted with either 2 ml or 1.3 ml 2X DNA/RNA Shield. Tissue samples were homogenized with an Omni TH homogenizer (Omni International). Prior to lysis the supernatants were divided into either four aliquots each containing 900 µl, or 3 aliquots each containing 950 µl, for NHP or human samples, respectively. The tissues were lysed by ballistic disintegration using a FastPrep-96 instrument (MP Biomedicals) and Zymo tubes containing 0.1 and 0.5-mm ZR BashingBeads (Zymo Research). Lysis was repeated three times at 1,600 rpm for 1 min, and tubes were placed on ice for 1 min after each lysis step. Particulate matter present in the supernatants was eliminated with QIAshredder homogenizers (Qiagen). RNA was isolated using the Zymobiomics RNA kit (Zymo Research) following the manufacturer instructions with the exceptions that all aliquots from a particular sample were pooled together before passing them through the first column and the recommended DNase treatment was performed twice for the human samples. Total RNA quality was assessed with an RNA Nanochip and an Agilent 2100 Bioanalyzer (Agilent Technologies).

### qRT-PCR analysis of infected NHP and human muscle

Total RNA extracted from infected NHP or human skeletal muscle was converted into cDNA using Superscript III reverse transcriptase, random primers, RNase OUT and dNTPs (all from Invitrogen). Quantitative RT-PCR (qRT-PCR) was performed using Taqman fast universal PCR master mix (Applied Biosystems) with an ABI 7500 Fast System instrument (Life Technologies). The genes tested were *Spy0014*, *Spy0271*, *braB*, *Spy1726*, *pptA*, and *glgP*. The sequences of primers and probes used in the qRT-PCR experiments are listed in the Table S9. Each experiment was performed with three technical replicates at three different dilutions. Transcript levels were normalized relative to the *rpsL* gene (encoding 30S ribosomal protein S12).

### Statistics

Results of lesion volume and CFU recovery from NHPs are expressed as mean ± SEM, with statistically significant differences determined using the Mann-Whitney test (Prism 6, Graphpad Software). Results of histology scoring of infected NHP muscle are expressed as mean ± SEM, with statistically significant differences determined using the Wilcoxon Rank Sum Test (Prism 6). Nonparametric tests were used because the data were shown to not follow a normal distribution using the Shapiro-Wilk test (Prism 6).

### Study approvals

All animal experiments were approved by the Institutional Animal Care and Use Committee of Houston Methodist Research Institute (protocol AUP-1217-0058). The human tissue was collected as part of a study approved by the Institutional Review Board at Houston Methodist Research Institute (protocol 0907-0151).

## Author contributions

LZ performed and analyzed TraDIS experiments, constructed and characterized isogenic mutant strains and wrote the manuscript. RJO planned and conducted experiments involving the NHPs, analyzed resulting data and wrote the manuscript. SBB analyzed the genomic data and wrote the manuscript. JME contributed critical discussions about transporter physiology and performed the TaqMan qRT-PCR to measure the transcript level of GAS transporter genes in infected NHP and human muscle. MOS performed the genome sequencing of the isogenic mutant strains and provided technical support for the NHP studies. SLK constructed and characterized isogenic mutant strains. CCC provided extensive technical support for all phases of the study. ARLC and ASW provided intellectual guidance for the TraDIS data analysis. LJ oversaw and performed the NHP experiments. JMM designed the studies, analyzed experiments, wrote the manuscript and oversaw the project.

## Supplemental material

## Acknowledgments

This work was supported by funds from the Fondren Foundation (to James M. Musser). Amelia R. L. Charbonneau is supported by the University of Cambridge Doctoral Training Partnership scheme, which is funded by the Biotechnology and Biological Sciences Research Council, UK (reference 1503883). We thank Drs. Frank DeLeo, Magnus Gottsfredsson, Karl G. Kristinsson, David M. Morens, and Kathryn E. Stockbauer for critical reading of the manuscript and suggesting improvements. We are indebted to Dr. Lillian S. Kao for providing tissue specimens from a patient with necrotizing myositis. We thank Annessa Smith and Caroline White for superb veterinary technical assistance.

## Conflicts of interest

The authors have declared that no conflicts of interest exist.

## Reference

1. Olsen RJ, and Musser JM. Molecular pathogenesis of necrotizing fasciitis. Annu Rev Pathol. 2010;5(1–31.

2. Stevens DL, and Bryant AE. Necrotizing Soft-Tissue Infections. N Engl J Med. 2017;377(23):2253–65.

3. Misiakos EP, Bagias G, Patapis P, Sotiropoulos D, Kanavidis P, and Machairas A. Current concepts in the management of necrotizing fasciitis. Front Surg. 2014;1(36.

4. Nelson GE, Pondo T, Toews KA, Farley MM, Lindegren ML, Lynfield R, Aragon D, Zansky SM, Watt JP, Cieslak PR, et al. Epidemiology Of Invasive Group A Streptococcal Infections in the United States, 2005-2012. Clin Infect Dis. 2016;63(4):478–86.

5. Lepoutre A, Doloy A, Bidet P, Leblond A, Perrocheau A, Bingen E, Trieu-Cuot P, Bouvet A, Poyart C, Levy-Bruhl D, et al. Epidemiology of invasive Streptococcus pyogenes infections in France in 2007. J Clin Microbiol. 2011;49(12):4094–100.

6. Carapetis JR, Steer AC, Mulholland EK, and Weber M. The global burden of group A streptococcal diseases. Lancet Infect Dis. 2005;5(11):685–94.

7. Cunningham MW. Pathogenesis of group A streptococcal infections. Clin Microbiol Rev. 2000;13(3):470–511.

8. Shannon O, Hertzen E, Norrby-Teglund A, Morgelin M, Sjobring U, and Bjorck L. Severe streptococcal infection is associated with M protein-induced platelet activation and thrombus formation. Mol Microbiol. 2007;65(5):1147–57.

9. Pahlman LI, Morgelin M, Eckert J, Johansson L, Russell W, Riesbeck K, Soehnlein O, Lindbom L, Norrby-Teglund A, Schumann RR, et al. Streptococcal M protein: a multipotent and powerful inducer of inflammation. J Immunol. 2006;177(2):1221–8.

10. Olsen RJ, Sitkiewicz I, Ayeras AA, Gonulal VE, Cantu C, Beres SB, Green NM, Lei B, Humbird T, Greaver J, et al. Decreased necrotizing fasciitis capacity caused by a single nucleotide mutation that alters a multiple gene virulence axis. Proc Natl Acad Sci U S A. 2010;107(2):888–93.

11. Hytonen J, Haataja S, Gerlach D, Podbielski A, and Finne J. The SpeB virulence factor of Streptococcus pyogenes, a multifunctional secreted and cell surface molecule with strepadhesin, laminin-binding and cysteine protease activity. Mol Microbiol. 2001;39(2):512–9.

12. Lukomski S, Montgomery CA, Rurangirwa J, Geske RS, Barrish JP, Adams GJ, and Musser JM. Extracellular cysteine protease produced by Streptococcus pyogenes participates in the pathogenesis of invasive skin infection and dissemination in mice. Infect Immun. 1999;67(4):1779–88.

13. Lukomski S, Burns EH Jr.,, Wyde PR, Podbielski A, Rurangirwa J, Moore-Poveda DK, and Musser JM. Genetic inactivation of an extracellular cysteine protease (SpeB) expressed by Streptococcus pyogenes decreases resistance to phagocytosis and dissemination to organs. Infect Immun. 1998;66(2):771–6.

14. Wessels MR, Moses AE, Goldberg JB, and DiCesare TJ. Hyaluronic acid capsule is a virulence factor for mucoid group A streptococci. Proc Natl Acad Sci U S A. 1991;88(19):8317–21.

15. Bricker AL, Carey VJ, and Wessels MR. Role of NADase in virulence in experimental invasive group A streptococcal infection. Infect Immun. 2005;73(10):6562–6.

16. Timmer AM, Timmer JC, Pence MA, Hsu LC, Ghochani M, Frey TG, Karin M, Salvesen GS, and Nizet V. Streptolysin O promotes group A Streptococcus immune evasion by accelerated macrophage apoptosis. J Biol Chem. 2009;284(2):862–71.

17. Zhu L, Olsen RJ, Lee JD, Porter AR, DeLeo FR, and Musser JM. Contribution of Secreted NADase and Streptolysin O to the Pathogenesis of Epidemic Serotype M1 Streptococcus pyogenes Infections. Am J Pathol. 2016.

18. Zhu L, Olsen RJ, Nasser W, Beres SB, Vuopio J, Kristinsson KG, Gottfredsson M, Porter AR, DeLeo FR, and Musser JM. A molecular trigger for intercontinental epidemics of group A Streptococcus. J Clin Invest. 2015;125(9):3545–59.

19. Zhu L, Olsen RJ, Nasser W, de la Riva Morales I, and Musser JM. Trading Capsule for Increased Cytotoxin Production: Contribution to Virulence of a Newly Emerged Clade of emm89 Streptococcus pyogenes. MBio. 2015;6(5):e01378–15.

20. Nasser W, Beres SB, Olsen RJ, Dean MA, Rice KA, Long SW, Kristinsson KG, Gottfredsson M, Vuopio J, Raisanen K, et al. Evolutionary pathway to increased virulence and epidemic group A Streptococcus disease derived from 3,615 genome sequences. Proc Natl Acad Sci U S A. 2014;111(17):E1768–76.

21. Beres SB, Kachroo P, Nasser W, Olsen RJ, Zhu L, Flores AR, de la Riva I, Paez-Mayorga J, Jimenez FE, Cantu C, et al. Transcriptome Remodeling Contributes to Epidemic Disease Caused by the Human Pathogen Streptococcus pyogenes. MBio. 2016;7(3).

22. Maruyama F, Watanabe T, and Nakagawa I. In: Ferretti JJ, Stevens DL, and Fischetti VA eds. Streptococcus pyogenes: Basic Biology to Clinical Manifestations. Oklahoma City (OK); 2016.

23. Beres SB, and Musser JM. Contribution of exogenous genetic elements to the group A Streptococcus metagenome. PLoS One. 2007;2(8):e800.

24. Kizy AE, and Neely MN. First Streptococcus pyogenes signature-tagged mutagenesis screen identifies novel virulence determinants. Infect Immun. 2009;77(5):1854–65.

25. Armbruster CE, Forsyth-DeOrnellas V, Johnson AO, Smith SN, Zhao L, Wu W, and Mobley HLT. Genome-wide transposon mutagenesis of Proteus mirabilis: Essential genes, fitness factors for catheter-associated urinary tract infection, and the impact of polymicrobial infection on fitness requirements. PLoS Pathog. 2017;13(6):e1006434.

26. Gawronski JD, Wong SM, Giannoukos G, Ward DV, and Akerley BJ. Tracking insertion mutants within libraries by deep sequencing and a genome-wide screen for Haemophilus genes required in the lung. Proc Natl Acad Sci U S A. 2009;106(38):16422–7.

27. Subashchandrabose S, Smith SN, Spurbeck RR, Kole MM, and Mobley HL. Genome-wide detection of fitness genes in uropathogenic Escherichia coli during systemic infection. PLoS Pathog. 2013;9(12):e1003788.

28. Wang N, Ozer EA, Mandel MJ, and Hauser AR. Genome-wide identification of Acinetobacter baumannii genes necessary for persistence in the lung. MBio. 2014;5(3):e01163–14.

29. Weerdenburg EM, Abdallah AM, Rangkuti F, Abd El Ghany M, Otto TD, Adroub SA, Molenaar D, Ummels R, Ter Veen K, van Stempvoort G, et al. Genome-wide transposon mutagenesis indicates that Mycobacterium marinum customizes its virulence mechanisms for survival and replication in different hosts. Infect Immun. 2015;83(5):1778–88.

30. Le Breton Y, Mistry P, Valdes KM, Quigley J, Kumar N, Tettelin H, and McIver KS. Genome-wide identification of genes required for fitness of group A Streptococcus in human blood. Infect Immun. 2013;81(3):862–75.

31. Zhu L, Charbonneau ARL, Waller AS, Olsen RJ, Beres SB, and Musser JM. Novel Genes Required for the Fitness of Streptococcus pyogenes in Human Saliva. mSphere. 2017;2(6).

32. Le Breton Y, Belew AT, Freiberg JA, Sundar GS, Islam E, Lieberman J, Shirtliff ME, Tettelin H, El-Sayed NM, and McIver KS. Genome-wide discovery of novel M1T1 group A streptococcal determinants important for fitness and virulence during soft-tissue infection. PLoS Pathog. 2017;13(8):e1006584.

33. Eraso JM, Olsen RJ, Beres SB, Kachroo P, Porter AR, Nasser W, Bernard PE, DeLeo FR, and Musser JM. Genomic Landscape of Intrahost Variation in Group A Streptococcus: Repeated and Abundant Mutational Inactivation of the fabT Gene Encoding a Regulator of Fatty Acid Synthesis. Infect Immun. 2016;84(12):3268–81.

34. Fittipaldi N, Beres SB, Olsen RJ, Kapur V, Shea PR, Watkins ME, Cantu CC, Laucirica DR, Jenkins L, Flores AR, et al. Full-genome dissection of an epidemic of severe invasive disease caused by a hypervirulent, recently emerged clone of group A Streptococcus. Am J Pathol. 2012;180(4):1522–34.

35. Sun H, Ringdahl U, Homeister JW, Fay WP, Engleberg NC, Yang AY, Rozek LS, Wang X, Sjobring U, and Ginsburg D. Plasminogen is a critical host pathogenicity factor for group A streptococcal infection. Science. 2004;305(5688):1283–6.

36. Sun H, Wang X, Degen JL, and Ginsburg D. Reduced thrombin generation increases host susceptibility to group A streptococcal infection. Blood. 2009;113(6):1358–64.

37. Kasper KJ, Zeppa JJ, Wakabayashi AT, Xu SX, Mazzuca DM, Welch I, Baroja ML, Kotb M, Cairns E, Cleary PP, et al. Bacterial superantigens promote acute nasopharyngeal infection by Streptococcus pyogenes in a human MHC Class II-dependent manner. PLoS Pathog. 2014;10(5):e1004155.

38. Sriskandan S, Unnikrishnan M, Krausz T, Dewchand H, Van Noorden S, Cohen J, and Altmann DM. Enhanced susceptibility to superantigen-associated streptococcal sepsis in human leukocyte antigen-DQ transgenic mice. J Infect Dis. 2001;184(2):166–73.

39. Marcum JA, and Kline DL. Species specificity of streptokinase. Comp Biochem Physiol B. 1983;75(3):389–94.

40. Wulf RJ, and Mertz ET. Studies on plasminogen. 8. Species specificity of streptokinase. Can J Biochem. 1969;47(10):927–31.

41. Langridge GC, Phan MD, Turner DJ, Perkins TT, Parts L, Haase J, Charles I, Maskell DJ, Peters SE, Dougan G, et al. Simultaneous assay of every Salmonella Typhi gene using one million transposon mutants. Genome Res. 2009;19(12):2308–16.

42. van Opijnen T, and Camilli A. Transposon insertion sequencing: a new tool for systems-level analysis of microorganisms. Nat Rev Microbiol. 2013;11(7):435–42.

43. Naseer U, Steinbakk M, Blystad H, and Caugant DA. Epidemiology of invasive group A streptococcal infections in Norway 2010-2014: A retrospective cohort study. Eur J Clin Microbiol Infect Dis. 2016;35(10):1639–48.

44. Smit PW, Lindholm L, Lyytikainen O, Jalava J, Patari-Sampo A, and Vuopio J. Epidemiology and emm types of invasive group A streptococcal infections in Finland, 2008-2013. Eur J Clin Microbiol Infect Dis. 2015;34(10):2131–6.

45. Meehan M, Murchan S, Gavin PJ, Drew RJ, and Cunney R. Epidemiology of an upsurge of invasive group A streptococcal infections in Ireland, 2012-2015. J Infect. 2018;77(3):183–90.

46. Smeesters PR, Laho D, Beall B, Steer AC, and Van Beneden CA. Seasonal, Geographic, and Temporal Trends of emm Clusters Associated With Invasive Group A Streptococcal Infections in US Multistate Surveillance. Clin Infect Dis. 2017;64(5):694–5.

47. Sumby P, Porcella SF, Madrigal AG, Barbian KD, Virtaneva K, Ricklefs SM, Sturdevant DE, Graham MR, Vuopio-Varkila J, Hoe NP, et al. Evolutionary origin and emergence of a highly successful clone of serotype M1 group a Streptococcus involved multiple horizontal gene transfer events. J Infect Dis. 2005;192(5):771–82.

48. Kachroo et al. Integrated analysis of population genomics, transcriptomics and virulence provides novel insights into Streptococcus pyogenes pathogenesis. In review.

49. Charbonneau ARL, Forman OP, Cain AK, Newland G, Robinson C, Boursnell M, Parkhill J, Leigh JA, Maskell DJ, and Waller AS. Defining the ABC of gene essentiality in streptococci. BMC Genomics. 2017;18(1):426.

50. Abel S, Abel zur Wiesch P, Davis BM, and Waldor MK. Analysis of Bottlenecks in Experimental Models of Infection. PLoS Pathog. 2015;11(6):e1004823.

51. Le Breton Y, Belew AT, Valdes KM, Islam E, Curry P, Tettelin H, Shirtliff ME, El-Sayed NM, and McIver KS. Essential Genes in the Core Genome of the Human Pathogen Streptococcus pyogenes. Sci Rep. 2015;5(9838.

52. Makthal N, Nguyen K, Do H, Gavagan M, Chandrangsu P, Helmann JD, Olsen RJ, and Kumaraswami M. A Critical Role of Zinc Importer AdcABC in Group A Streptococcus-Host Interactions During Infection and Its Implications for Vaccine Development. EBioMedicine. 2017;21(131–41.

53. Sanson M, Makthal N, Flores AR, Olsen RJ, Musser JM, and Kumaraswami M. Adhesin competence repressor (AdcR) from Streptococcus pyogenes controls adaptive responses to zinc limitation and contributes to virulence. Nucleic Acids Res. 2015;43(1):418–32.

54. van Sorge NM, Cole JN, Kuipers K, Henningham A, Aziz RK, Kasirer-Friede A, Lin L, Berends ETM, Davies MR, Dougan G, et al. The classical lancefield antigen of group a Streptococcus is a virulence determinant with implications for vaccine design. Cell Host Microbe. 2014;15(6):729–40.

55. Henningham A, Davies MR, Uchiyama S, van Sorge NM, Lund S, Chen KT, Walker MJ, Cole JN, and Nizet V. Virulence Role of the GlcNAc Side Chain of the Lancefield Cell Wall Carbohydrate Antigen in Non-M1-Serotype Group A Streptococcus. MBio. 2018;9(1).

56. Honda-Ogawa M, Sumitomo T, Mori Y, Hamd DT, Ogawa T, Yamaguchi M, Nakata M, and Kawabata S. Streptococcus pyogenes Endopeptidase O Contributes to Evasion from Complement-mediated Bacteriolysis via Binding to Human Complement Factor C1q. J Biol Chem. 2017;292(10):4244–54.

57. Brouwer S, Cork AJ, Ong CY, Barnett TC, West NP, McIver KS, and Walker MJ. Endopeptidase PepO Regulates the SpeB Cysteine Protease and Is Essential for the Virulence of Invasive M1T1 Streptococcus pyogenes. J Bacteriol. 2018;200(8).

58. Reid SD, Montgomery AG, Voyich JM, DeLeo FR, Lei B, Ireland RM, Green NM, Liu M, Lukomski S, and Musser JM. Characterization of an extracellular virulence factor made by group A Streptococcus with homology to the Listeria monocytogenes internalin family of proteins. Infect Immun. 2003;71(12):7043–52.

59. Brenot A, King KY, and Caparon MG. The PerR regulon in peroxide resistance and virulence of Streptococcus pyogenes. Mol Microbiol. 2005;55(1):221–34.

60. Trevino J, Liu Z, Cao TN, Ramirez-Pena E, and Sumby P. RivR is a negative regulator of virulence factor expression in group A Streptococcus. Infect Immun. 2013;81(1):364–72.

61. Lynskey NN, Goulding D, Gierula M, Turner CE, Dougan G, Edwards RJ, and Sriskandan S. RocA truncation underpins hyper-encapsulation, carriage longevity and transmissibility of serotype M18 group A streptococci. PLoS Pathog. 2013;9(12):e1003842.

62. Jain I, Miller EW, Danger JL, Pflughoeft KJ, and Sumby P. RocA Is an Accessory Protein to the Virulence-Regulating CovRS Two-Component System in Group A Streptococcus. Infect Immun. 2017;85(11).

63. Biswas I, and Scott JR. Identification of rocA, a positive regulator of covR expression in the group A streptococcus. J Bacteriol. 2003;185(10):3081–90.

64. Miller EW, Danger JL, Ramalinga AB, Horstmann N, Shelburne SA, and Sumby P. Regulatory rewiring confers serotype-specific hyper-virulence in the human pathogen group A Streptococcus. Mol Microbiol. 2015;98(3):473–89.

65. Zhu L, Olsen RJ, Horstmann N, Shelburne SA, Fan J, Hu Y, and Musser JM. Intergenic Variable-Number Tandem-Repeat Polymorphism Upstream of rocA Alters Toxin Production and Enhances Virulence in Streptococcus pyogenes. Infect Immun. 2016;84(7):2086–93.

66. Bernard PE, Kachroo P, Zhu L, Beres SB, Eraso JM, Kajani Z, Long SW, Musser JM, and Olsen RJ. RocA has serotype-specific gene regulatory and pathogenesis activity in serotype M28 group A streptococcus. Infect Immun. 2018.

67. Basavanna S, Chimalapati S, Maqbool A, Rubbo B, Yuste J, Wilson RJ, Hosie A, Ogunniyi AD, Paton JC, Thomas G, et al. The effects of methionine acquisition and synthesis on Streptococcus pneumoniae growth and virulence. PLoS One. 2013;8(1):e49638.

68. Davies HC, Karush F, and Rudd JH. Effect of Amino Acids on Steady-State Growth of a Group a Hemolytic Streptococcus. J Bacteriol. 1965;89(421–7.

69. Pancholi V, and Caparon M. In: Ferretti JJ, Stevens DL, and Fischetti VA eds. Streptococcus pyogenes: Basic Biology to Clinical Manifestations. Oklahoma City (OK); 2016.

70. Rodriguez-Ortega MJ, Norais N, Bensi G, Liberatori S, Capo S, Mora M, Scarselli M, Doro F, Ferrari G, Garaguso I, et al. Characterization and identification of vaccine candidate proteins through analysis of the group A Streptococcus surface proteome. Nat Biotechnol. 2006;24(2):191–7.

71. Chang JC, and Federle MJ. PptAB Exports Rgg Quorum-Sensing Peptides in Streptococcus. PLoS One. 2016;11(12):e0168461.

72. Alonso-Casajus N, Dauvillee D, Viale AM, Munoz FJ, Baroja-Fernandez E, Moran-Zorzano MT, Eydallin G, Ball S, and Pozueta-Romero J. Glycogen phosphorylase, the product of the glgP Gene, catalyzes glycogen breakdown by removing glucose units from the nonreducing ends in Escherichia coli. J Bacteriol. 2006;188(14):5266–72.

73. Chimalapati S, Cohen JM, Camberlein E, MacDonald N, Durmort C, Vernet T, Hermans PW, Mitchell T, and Brown JS. Effects of deletion of the Streptococcus pneumoniae lipoprotein diacylglyceryl transferase gene lgt on ABC transporter function and on growth in vivo. PLoS One. 2012;7(7):e41393.

74. Das S, Kanamoto T, Ge X, Xu P, Unoki T, Munro CL, and Kitten T. Contribution of lipoproteins and lipoprotein processing to endocarditis virulence in Streptococcus sanguinis. J Bacteriol. 2009;191(13):4166–79.

75. Weston BF, Brenot A, and Caparon MG. The metal homeostasis protein, Lsp, of Streptococcus pyogenes is necessary for acquisition of zinc and virulence. Infect Immun. 2009;77(7):2840–8.

76. Voyich JM, Braughton KR, Sturdevant DE, Vuong C, Kobayashi SD, Porcella SF, Otto M, Musser JM, and DeLeo FR. Engagement of the pathogen survival response used by group A Streptococcus to avert destruction by innate host defense. J Immunol. 2004;173(2):1194–201.

77. Voyich JM, Sturdevant DE, Braughton KR, Kobayashi SD, Lei B, Virtaneva K, Dorward DW, Musser JM, and DeLeo FR. Genome-wide protective response used by group A Streptococcus to evade destruction by human polymorphonuclear leukocytes. Proc Natl Acad Sci U S A. 2003;100(4):1996–2001.

78. Bergstrom J, Furst P, Noree LO, and Vinnars E. Intracellular free amino acid concentration in human muscle tissue. J Appl Physiol. 1974;36(6):693–7.

79. Chang JC, LaSarre B, Jimenez JC, Aggarwal C, and Federle MJ. Two group A streptococcal peptide pheromones act through opposing Rgg regulators to control biofilm development. PLoS Pathog. 2011;7(8):e1002190.

80. Graham MR, Virtaneva K, Porcella SF, Barry WT, Gowen BB, Johnson CR, Wright FA, and Musser JM. Group A Streptococcus transcriptome dynamics during growth in human blood reveals bacterial adaptive and survival strategies. Am J Pathol. 2005;166(2):455–65.

81. Hertzen E, Johansson L, Kansal R, Hecht A, Dahesh S, Janos M, Nizet V, Kotb M, and Norrby-Teglund A. Intracellular Streptococcus pyogenes in human macrophages display an altered gene expression profile. PLoS One. 2012;7(4):e35218.

82. Graham MR, Smoot LM, Migliaccio CA, Virtaneva K, Sturdevant DE, Porcella SF, Federle MJ, Adams GJ, Scott JR, and Musser JM. Virulence control in group A Streptococcus by a two-component gene regulatory system: global expression profiling and in vivo infection modeling. Proc Natl Acad Sci U S A. 2002;99(21):13855–60.

83. Barquist L, Mayho M, Cummins C, Cain AK, Boinett CJ, Page AJ, Langridge GC, Quail MA, Keane JA, and Parkhill J. The TraDIS toolkit: sequencing and analysis for dense transposon mutant libraries. Bioinformatics. 2016;32(7):1109–11.

